# Enhanced Detection of Glioblastoma Vasculature with Superparamagnetic Iron Oxide Nanoparticles and MRI

**DOI:** 10.1101/2025.02.02.635583

**Authors:** Phillip W. Janowicz, Thomas Boele, Richard T. Maschmeyer, Yaser H. Gholami, Emma G. Kempe, Brett W. Stringer, Shihani P. Stoner, Marie Zhang, Taymin du-Toit Thompson, Fern Williams, Aude Touffu, Lenka Munoz, Zdenka Kuncic, Caterina Brighi, David E. J. Waddington

## Abstract

Detecting glioblastoma infiltration in the brain is challenging due to limited MRI contrast beyond the enhancing tumour core. This study aims to investigate the potential of superparamagnetic iron oxide nanoparticles (SPIONs) as contrast agents for improved detection of diffuse brain cancer. We examine the distribution and pharmacokinetics of SPIONs in glioblastoma models with intact and disrupted blood-brain barriers. Using MRI, we imaged RN1-luc and U87MG mice injected with Gadovist and SPIONs, observing differences in blood-brain barrier permeability. Peripheral imaging showed strong uptake of nanoparticles in the liver and spleen, while vascular and renal signals were transient. Susceptibility gradient mapping enabled positive nanoparticle contrast within tumours and provided additional information on tumour angiogenesis. This approach offers a novel method for detecting diffuse brain cancer. Our findings demonstrate that SPIONs enhance glioblastoma detection beyond conventional MRI, providing insights into tumour angiogenesis and opening new avenues for early diagnosis and targeted treatment strategies.

## Introduction

Glioblastoma is the most common type of malignant brain cancer and its prognosis is poor^1^. A significant challenge to the treatment of glioblastoma is the heterogenous and invasive nature of the tumour, which leads to inconsistent disruption of the blood brain barrier (BBB). While the BBB is typically disrupted within the tumour core, in regions of tumour infiltration, the BBB remains intact despite the presence of clinically significant tumour burden^2^. As most diagnostic contrast agents and cancer therapeutics do not cross an intact BBB, regions of infiltrating tumour are extremely difficult to both identify and treat^3^. In particular, while magnetic resonance imaging (MRI) of gadolinium-based contrast agents (GBCAs) is part of the standard of care for glioblastoma diagnosis, GBCAs such as Gadovist only highlight tumour in regions of significant BBB disruption^4^.

Conventional imaging of glioblastoma consists of T1-weighted (T1w) and T2-weighted (T2w) scans in addition to a T1-contrast enhanced (T1-CE) scan acquired after intravenous administration of a GBCA (Figure 1A). While T1-CE scans boost signal in the core tumour mass, regions of hyperintensity in T2w scans beyond this core mass are a mix of infiltrative tumour lying behind an intact BBB and oedema of normal tissue^5^. Distinguishing between tumour and normal tissue is key to surgical and radiotherapy treatment planning. Hence, essential to the improved diagnosis and treatment of glioblastoma is the development of diagnostic tools that can provide information on tumour growth despite the presence of an intact BBB^6^.

**Figure 1:**
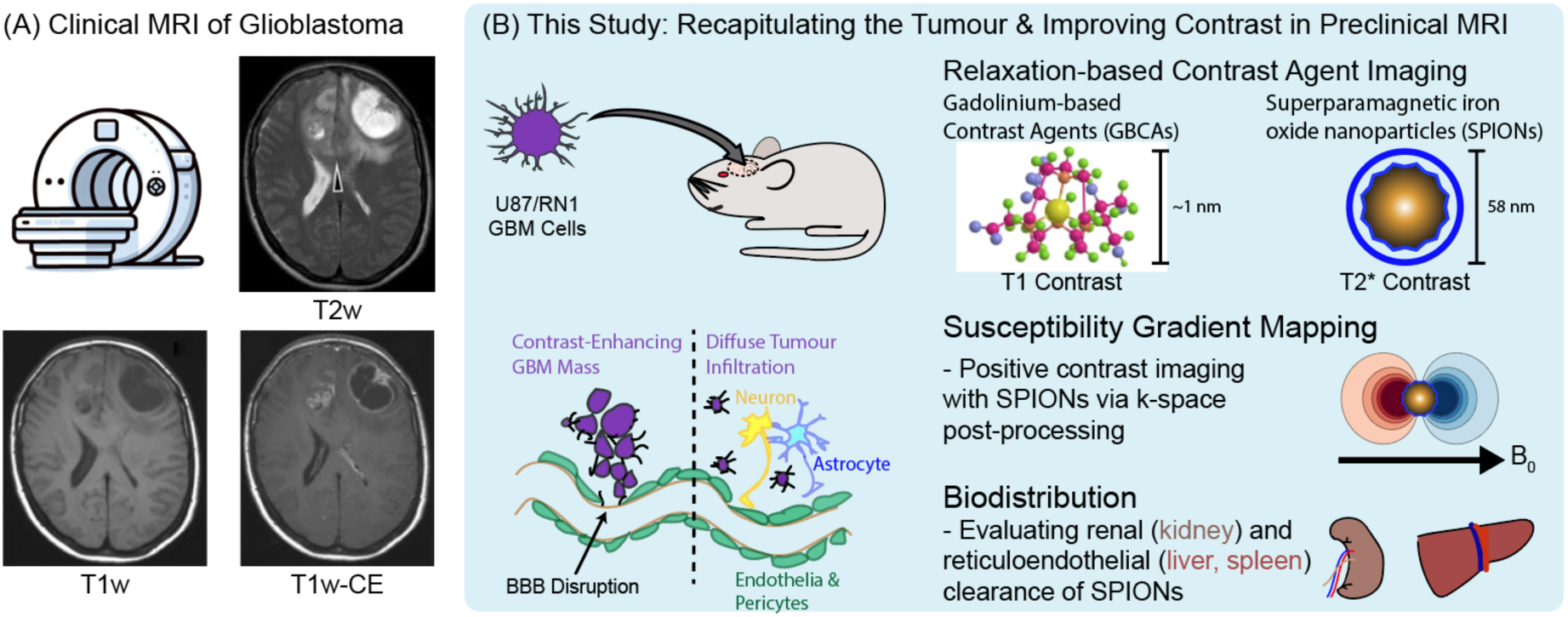
**A)** Clinical example of mismatch between glioblastoma T2 hyperintensity and areas of T1 gadolinium contrast enhancement. **B)** To replicate the heterogeneously disrupted blood brain barrier (BBB) in clinical glioblastoma (GBM), we used a diffuse, infiltrative patient-derived xenograft, RN1, to compare PEG-SPION to Gadovist in glioblastoma, as well as in non-diffuse glioblastoma U87. We also tested the effect of PEGylation on the SPION biodistribution and generated positive SPION contrast using susceptibility gradient mapping. Human data was adapted from Claes, et al. ^27^, with open access permission from Springer (all rights reserved).

MRI contrast agents based on nanoparticles are of increasing interest for their ability to probe additional factors within the tumour environment and could help to differentiate infiltrative tumour and healthy tissues in MRI scans. In particular, superparamagnetic iron oxide nanoparticles (SPIONs), which are nanoparticles with a 3-25 nm core of iron oxide (typically magnetite Fe_3_O_4_ or maghemite Fe_2_O_3_), are of interest for their significant MRI contrast potential and biocompatibility^7–9^. SPIONs enhance MRI contrast by shortening transverse relaxation times of nearby protons (T2 contrast) and by creating strong nanoscale magnetic inhomogeneities (T2* contrast)^10–12^. As a highly sensitive contrast agent, with long blood circulation times, there is significant potential for SPIONs to be a complementary contrast method that distinguishes regions of infiltrative tumour using MRI.

Additionally, there are growing concerns over the accumulation of GBCAs in central-nervous-system tissue, even in patients with normal renal function, which may constrain the use of GBCAs into the future^13–21^. Due to their larger size, SPIONs bypass renal deposition, and are metabolised into bioavailable haemoglobin^22^ rather than being deposited enduringly in tissue. Despite the improved safety profile of SPIONs, T2 relaxivity mechanisms lead to negative MRI contrast (reduced signal). Negative contrast is typically disliked by radiologists^23^ and there have been limited efforts to substitute SPIONs for GBCAs, which give positive contrast in T1w MRI.

Here, we preclinically evaluate the use of SPIONs for identification of diffuse glioblastoma with MRI. First, we recapitulate glioblastoma in a mouse model (Figure 1B) using cell lines known to grow aggressive tumours with a highly disrupted BBB (U87MG, GBM6) and a cell line that grows infiltrative tumours with an intact BBB (RN1)^24,25^. Second, we compare conventional MRI approaches to imaging glioblastoma to MRI scans acquired with SPIONs, showing that we can image angiogenesis within infiltrative tumour that is not detected with GBCAs. Third, we implement a postprocessing method called susceptibility gradient mapping (SGM)^26^ that generates ‘positive’ contrast rather than ‘negative’ MRI contrast from T2*w scans with SPIONs, overcoming one of the main barriers to the clinical use of SPIONs. We conclude by evaluating the biodistribution of SPIONs, showing that nanoparticle clearance predominantly occurs via the reticuloendothelial system. Our results show the potential of SPIONs to enhance the imaging of glioblastoma with MRI and will inform the development of new approaches to imaging and treating this disease in the clinic.

## Methods

### Materials

The nanoparticles evaluated in this study are novel magnetite(Fe_3_O_4_)-based, methoxy-PEG-coated SPIONs^28^. PEG-SPIONs (mPEG PrecisionMRX) were sourced commercially and received in aqueous solutions with elemental iron concentrations ranging from 89 to 374 mM (~5 to 20.9 mg/mL). PEG-SPIONs have a hydrodynamic diameter of 58.3 nm (measured by dynamic light scattering), core diameter of 24.2 nm ±1.3 nm (measured by Rigaku SmartLab small angle X-ray scattering and JEOL 1200EX transmission electron microscopy) and polydispersity index of 0.06. Originally designed for magnetic relaxometry^29^, which operates at mT field strengths, these mPEG-SPIONs were shown to generate *in vivo* contrast in ultra-low field MRI (6.5 mT)^30,31^.

In addition to brain tumour imaging with PEG-SPIONs, we also evaluated the biodistribution of PEG-SPIONs and compare this to Ferumoxytol. Ferumoxytol is a SPION composed of a magnetite core of 3.25 nm diameter, which is coated with polyglucose sorbitol carboxymethylether, yielding nanoparticles with a hydrodynamic diameter of 26.3 nm. Both samples were diluted in either deoxygenated MilliQ water for relaxometric studies^30^ or in sterile saline for animal studies. All cell culturing reagents were supplied by Thermo Fisher Scientific unless otherwise specified.

### Cell Culture

All cell cultures were incubated at 37 °C in 5% CO_2_. RN1-luciferase (RN1-luc)^24^ and GBM6^32^ cells were grown for a maximum of five passages in KnockOut^TM^ DMEM/F-12 basal media (Gibco) supplemented with epidermal growth factor (20 ng/mL), basic fibroblast growth factor (20 ng/mL), 2% StemPro^TM^ neural supplement, GlutaMAX^TM^ supplement (2 mM), 100 U/mL penicillin and 100 µg/mL streptomycin on T75 flasks coated with 0.15% Matrigel (Sigma-Aldrich) in phosphate buffered saline (PBS). RN1 is a patient-derived xenograft isolated from a 56 year old male with a primary isocitrate dehydrogenase 1 (IDH1) wild-type, O-6-methylguanine-DNA methyltransferase (MGMT) unmethylated promoter glioblastoma located in the left temporal lobe^28^. RN1-luc cells were selected for using 1 µg/mL blasticidin S hydrochloride (Cayman Chemical). Accutase was used for cell passaging. U87MG cells were grown in DMEM supplemented with 10% foetal calf serum, 100 U/mL penicillin and 100 µg/mL streptomycin.

### Animals and Glioblastoma Engraftment Surgery

Sixteen female mice were included in this study, and were five weeks of age at the time of surgery. Nonobese diabetic Prkdc^SCID^ (NOD-SCID) mice were used for the brain cancer imaging component of the study. Due to a supply shortage of NOD-SCID mice, GBM6 tumours were grown in NODRag1^null^ IL2rγ^null^ (NRG) mice, which is a similar strain. Nanoparticle biodistribution was evaluated in an immunocompetent mouse strain (C57BL/6).

All experimental protocols and procedures were approved by the University of Sydney Animal Ethics Committee, under protocols 1316 (nanoparticle biodistribution) and 2170 (brain cancer). All methods are reported in accordance with the ARRIVE guidelines. All methods were carried out in accordance with local guidelines and regulations. Mice were monitored for weight changes and symptoms following the mouse grimace scale^33^ at least twice per week throughout the study period. Mice were euthanized via anaesthesia overdose, confirmed by cervical dislocation, at the conclusion of final imaging timepoints for each study.

For tumour engraftment, passaged cell pellets were resuspended in 10 ng/mL laminin from mouse Engelbreth-Holm-Swarm sarcoma (Merck) in PBS to give 100-150k cells per µL and they were kept on ice during the surgeries. Surgical tools and cotton bud tips were autoclaved prior to use. Each animal was assessed as per the mouse grimace scale for pain evaluation and monitored and weighed prior to surgery and intensively observed for the following three days. Mice were anaesthetised with 3% isoflurane and maintained under 2% isoflurane in a stereotaxic frame (Kopf Instruments) with ear bars, and breaths per minute were monitored by an assistant. Once deep anaesthesia was confirmed with a toe pinch, the scalp was shaved and washed with pHisohex (Viatris), and an approximate 10-15 mm long incision was made along the skin above the sagittal suture. A hole was drilled gently 0.1 mm posterior to bregma and 2 mm laterally, and 200k (U87MG), 250k (GBM6) or 300k (RN1-luc) cells were loaded into a 2 µL 25 G bevelled Hamilton syringe. The needle was inserted 3 mm into the brain slowly over a 2 min period, then retracted 0.5 mm and left to sit for 2 min. The cells were injected slowly at ~0.7 µL/min. At the conclusion of the injection, the needle was left in place for 2 min allowing the cells to settle, and retracted slowly. The burr hole was sealed with a small amount of bone wax (Ethicon) and the skin was held closed with forceps and glued using 3M Vetbond. An analgesic subcutaneous injection of 0.1 mg/kg buprenorphine (Buprelieve®, Jurox) and an anti-inflammatory second subcutaneous injection of 5 mg/kg meloxicam (Apex) were given before turning off the isoflurane and placing the mouse in a recovery cage with body temperature warming and post-operative monitoring.

### *In vivo* bioluminescence imaging

RN1-luc PDX progression was monitored fortnightly to weekly using either 7 T MRI with T2 weighted fast spin echo sequences (described below) or by bioluminescence using a Perkin Elmer *In Vivo* Imaging System (IVIS). Mice were anaesthetised using isoflurane, had scalps shaved and were injected intraperitoneally with 150 mg/kg VivoGlo^TM^ D-luciferin (Promega). Each mouse was imaged 5 min post-injection using standard bioluminescence filters as per the Living Image software.

### Magnetic Resonance Imaging

All MR imaging experiments were performed on MR Solutions preclinical MRI scanners. While the collection of data on a 7 T preclinical system was our preferred plan, due to unavoidable system outages, some data were collected on a 3 T preclinical system. For biodistribution experiments, three mice were imaged at 3 T and three at 7 T, while for glioblastoma brain imaging, four RN1-luc, two U87MG and three GBM6 mice were imaged at 7 T, and one U87MG mouse was imaged at 3 T.

#### Brain cancer imaging

Mice were transferred to our preclinical imaging facility after allowing 3 days for recovery from tumour inoculation. They were subsequently allowed 1 week acclimatization in the new facility before imaging could take place. Mice brain xenografts were monitored by MRI from two weeks post-engraftment for U87MG and GBM6, and from approximately 35 days post-engraftment for RN1-luc, onwards. MRI acquisitions to monitor tumour growth were performed which included T2-weighted fast spin echo sequences (repetition time (TR) = 4000 ms, echo time (TE) = 45 ms, flip angle (FA) = 90°, field of view (FOV) = 25×25 mm, number of averages (NA) = 3, and slice thickness (ST) = 0.5 mm) in the coronal plane as well as the sagittal and horizontal planes when necessary, using a mouse head coil in a 7 T MR Solutions scanner. Once tumours appeared sufficiently large (>2-3 mm diameter) on the T2-weighted scans, or IVIS, mice proceeded to the Gadovist and SPIONs injection imaging stage of the study. Mice were intravenously injected with 0.1 mmol/kg Gadovist, followed by 50 µL heparin sodium (50 injection units/mL, Hospira) in saline via cannula.

Mice were subsequently injected with SPIONs after all Gadovist had cleared from the brain (at least 20 minutes afterwards). GBCAs are known to rapidly clear from mouse circulation with a half-life of 5-6 minutes^34^. Anatomical scans including the above mentioned T2-weighted scans and also T1-weighted fast spin echo scans (TR = 1000 ms, TR = 11 ms, FA = 90°, FOV = 25×25 mm, NA = 4, SL = 0.5 mm) and variable flip angle T1 mapping sequences (data not shown) were acquired pre- and post-Gadovist injection. For SPIONs imaging, T2*-weighted fast low angle shot (FLASH) gradient recalled echo (GRE) sequences (TR = 400 ms, TE = 9 ms, FA = 40°, FOV = 25×25 mm, NA = 4, SL = 0.5 mm) and T2*-weighted multi-gradient echo sequences (TR = 500 ms, TE = 4, 8.48, 12.96, 17.44, 21.92 and 26.4 ms, FA = 50°, FOV = 25×25 mm, NA = 4, ST = 0.5 mm) were acquired pre- and post-injection. T2*-weighted images were acquired immediately after PEG-SPION injection and were repeated up to 1 h post SPION injection. Repeat imaging sessions were conducted at later timepoints of 2, 3, 5 and 24 h.

We note that there was no need for randomisation or blinding in our study design as all mice received Gadovist and SPION injections. All animals were included in the study and subsequent data analysis unless a brain tumour failed to develop after inoculation with a glioblastoma cell line. Due to the aggressive nature of the tumour model, the study was designed to minimise the number of mice used. A sample size of *n=3* was sufficient for distinguishing between tumour types and contrast effects.

#### Biodistribution imaging

C57BL/6 mice were used for the pharmacokinetics component of the study to evaluate biodistribution in an immunocompetent setting, using 3 T MRI. Mice were sedated with isoflurane and injected slowly over 1 min into the lateral tail vein via a cannula with 0.2 ml of saline containing 0.2-1.25 mg Fe/mL PEG-SPION or ferumoxytol, resulting in 2, 5 or 10 mg Fe/kg dose injected intravenously. For mice imaged at 7 T and 3 T for PEG-SPION pharmacokinetics, 5 and 10 mg/kg was injected respectively, whereas for ferumoxytol at 3 T, 2 mg/kg was injected. Mice were anaesthetised using 3% isoflurane and maintained with 1-2% isoflurane for the duration of the imaging period. For 3 T pharmacokinetics nanoparticle injections, mice were imaged at pre-injection (N), and 1, 6, 24, 48, 72 and 96 hours post-injection using fast spin echo T1-weighted (TR = 350 ms, TE = 11ms, ST = 1 mm, NA = 10, number of slices = 12), T2-weighted sequences (TR = 2600ms, TE = 72 ms, ST = 1 mm, NOA = 6, number of slices = 16), and GRE FLASH T2* weighted sequences (TR = 143 ms for PEG-SPION and TR = 240 ms for ferumoxytol, TE = 9 ms, NA = 10, ST = 1 mm, number of slices = 8), using a mouse body coil. In a latter separate study, NOD/SCID mice were injected with 5 mg/kg PEG-SPION and imaged on a 7T MR Scanner (MR Solutions) with similar sequences. The 7T sequences included a T1-weighted fast spin echo (TR = 1000 ms, TE = 11 ms, NA = 3, ST = 1 mm, FOV = 45×60 mm) and a T2*-weighted GRE FLASH (TR = 400 ms, TE= 9 ms, FA = 40°, number of averages = 4, slice thickness = 1 mm, field of view = 60×45 mm).

### Histology and Microscopy

Mice were transcardially perfused with 25 mL saline followed by 25 mL 10% neutral buffered formalin using a 25 G butterfly needle (Terumo) while under deep 3% isoflurane anaesthesia. Fixed tissue was transferred to 80% ethanol after 24-48 h immersion in 10% neutral buffered formalin (4 °C) and then dehydrated and paraffin embedded. Ten-micron sections were acquired onto Superfrost Plus slides using an Epredia rotary microtome with Feather S35 microtome blades including the brain, liver, spleen, heart, kidneys, and lung. Sections were stained with Perls’ Prussian blue using 2% potassium ferrocyanide mixed 1:1 with 2% hydrochloric acid immediately before use for 15 min, followed by 5 min 0.1% acetic safranin solution (0.1% glacial acetic acid, 0.1% safranin O) counterstain and cover-slipped under dibutylphthalate polystyrene xylene (DPX) mountant. Brightfield images were acquired using a Zeiss AxioScan Z1 slide scanner equipped with a Plan-Apochromat 20x objective and Hitachi HV-F202SCL camera, and visualised with ZEN v3.1 software.

### Image processing and analysis

SPION biodistribution analyses were performed by comparing the mean grey values (MGV) of regions of interest (ROIs) across MRI slices of key organs (brain, liver, spleen, kidneys, lung, stomach, leg and back muscle, and bladder) at timepoints from pre-injection to 1, 6, 24, 48, 72 and 96 hours post-injection. All T1, T2 and T2* weighted images were analysed. DICOMs produced by the MR Solutions preclinical MRI scanner are auto-scaled to the maximum signal intensity. Hence, for quantitative analyses, scaling-invariant images were reconstructed from raw k-space data using an inverse fast Fourier transform operation in MATLAB. For data analysis, DICOM figures were exported from MATLAB and ROIs were drawn using MicroDICOM v2024.2 software. The average ROI MGV across slices is presented in Supplementary Figure 3 with error bars representing the standard error of the mean, while in Figure 5C the average MGV of multiple slices across the two mice per timepoint was used. For Figure 3C, post-injection divided by pre-injection values were determined.

### Positive contrast post-processing using susceptibility gradient mapping

SGM is a positive contrast method that is not reliant on longitudinal relaxation like conventional T1 positive contrast agents, but rather makes use of the fact that T2* contrast agents create local susceptibility gradients that dephase precessing magnetic moments. SGM leverages the complex k-space data obtained from a conventional GRE sequence, calculates local magnetic field gradients associated with variations in magnetic susceptibility and visualizes them in a parametric map. The following equation^26^ defines the magnitude of the susceptibility gradient map in one dimension:

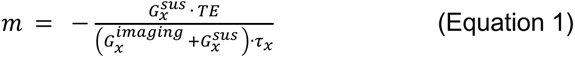

Here, *τ_x_* is the sampling rate, 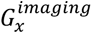 is the imaging gradient and 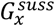 is the susceptibility gradient that causes an unbalanced timing of the gradient echo at *TE*.SGMs ^26^were produced from the raw *k*-space T2* gradient recalled echo data by performing short-term 1D Fourier transforms on each voxel and its two adjacent voxels following the procedure first described in Dahnke, et al. ^26^. A fit was performed to calculate the magnitude of the susceptibility gradient vector. This was performed in both the x and y directions, with the two results then being combined to produce an overall SGM. This process was applied to skull stripped images of the mouse brain so as to minimize the visual contribution of susceptibility artefacts at air-tissue interfaces, which are amplified by the SGM fitting procedure.

### Statistical Analysis

All data is presented as mean + standard error of the mean (SEM). Pre versus post-injection brain MGVs were compared using a two-way analysis of variance (ANOVA) with a Šídák’s multiple comparison test in GraphPad Prism 10.3 Software.

## Results

### RN1-luc xenografts have a Gadovist-impermeable blood brain barrier in contrast to U87MG xenografts

U87MG orthotopic xenografts appear highly permeable towards 0.1 mmol/kg Gadovist®, as shown by the positive contrast in T1-weighted turbo spin echo sequences post-Gadovist injection (Figure 2A). However, RN1-luc stem-like patient-derived orthotopic xenografts displayed limited contrast after Gadovist injection (Figure 2A), likely indicating a Gadovist impermeable BBB despite T2 hyperintensity. Tumour infiltration by RN1-luc in regions that did not show tumour infiltration was later confirmed by histology (Figure 2B). The limited BBB permeability of Gadovist observed in RN1-luc infiltrative tumour tissue was also observed in another patient-derived xenograft, GBM6 (Figure S1).

**Figure 2 –.**
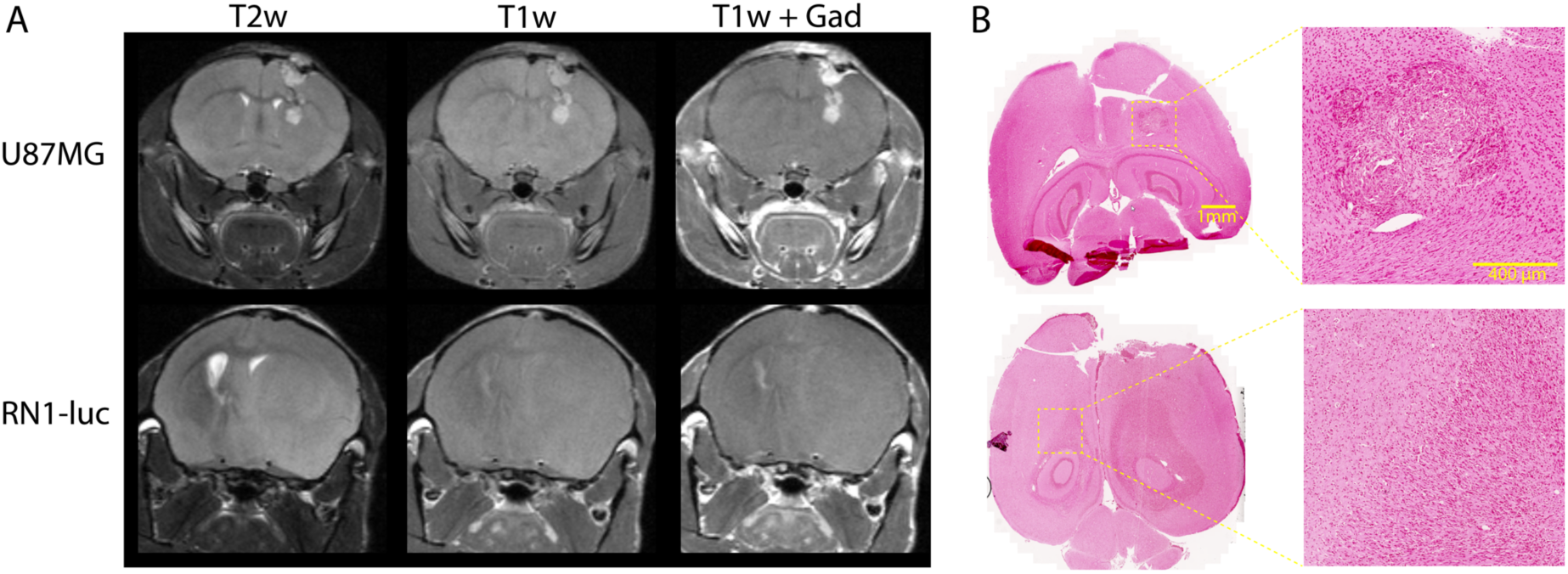
**A)** Representative examples of U87MG and RN1-luc xenograft anatomy in T1 and T2 weighted scans, including post-Gadovist (0.1 mmol/kg). **B)** Brain histology of U87MG and RN1-luc patient-derived orthotopic xenografts, stained with Perls’ Prussian blue and counterstained with 0.1% safranin O. RN1-luc shows a distinctly diffuse phenotype whereby it has spread laterally across hemispheres, whereas U87MG shows a demarcated non-diffuse phenotype. Scale bars = 1 mm and 400 microns respectively (images acquired using a 20x objective).

### Glioblastoma angiogenesis was contrast enhanced via SPION T2* vascular signal

Patient-derived orthotopic xenografts RN1-luc and U87MG were evaluated for PEG-SPION uptake using T2*, T2 and T1 weighted scans. In both tumours the nanoparticles showed strong T2* contrast effects, which were predominantly located in normal neurovasculature and angiogenic tumour areas (Figure 3). There was a statistically significant change in tumour vascular contrast in U87MG and RN1-luc compared to control regions following injection of PEG-SPION (Figure 3C). The PEG-SPION T2* contrast was largely washed out from the vasculature after 5 h from injection, with no detectable amount of signal in the RN1-luc tumour area at 24 h. However, both U87MG and RN1-luc tumour vascular PEG-SPION T2* contrast was visible at 2 h and 3 h post injection. The lack of Perls’ Prussian blue staining in the tumour area 1 h post-perfusion (Figure 2B) is indicative that the PEG-SPION was localised to the tumour vasculature rather than tumour cells themselves.

**Figure 3 –.**
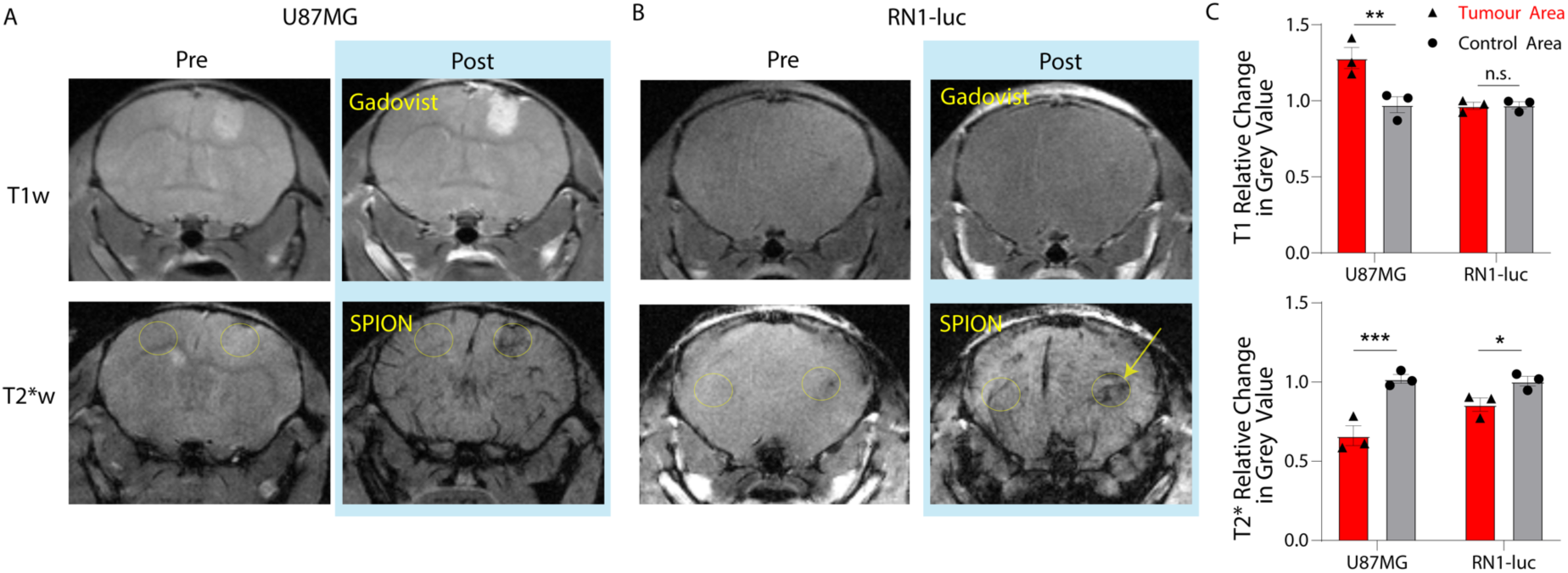
T1- and T2*-weighted MR images of **(A)** U87MG and **(B)** RN1-luc patient-derived xenografts pre- and two minutes post-injection of either 0.1 mmol/kg Gadovist or 10 mg/kg PEG-SPIONs. **C)** Quantification of change in pre- to post-injection from representative regions of interest (ROIs) in T1 and T2* weighted images. ROIs were drawn around the brighter (T1w) or darker (T2*w, e.g. yellow arrow) appearing tumour regions and quantified relative to the same ROIs in the pre-injection images, and compared to control hemisphere non-tumour area, giving mean ± SEM post/pre injection values (n=3). * denotes p<0.05, ** p<0.01 and *** p<0.001, as measured by two-way ANOVA with Šídák’s multiple comparisons test.

### SPION Positive Contrast from Quantitative Susceptibility Gradient Mapping

We found that SGM can highlight PEG-SPIONs in the U87MG and RN1-luc tumours as positive contrast, with a trade-off being a lower effective resolution (Figure 4). In the case of RN1-luc which has a functional BBB, the borders of this tumour were not clear from T2* SGM but instead abnormal vasculature and blood pooling in the tumour area appeared bright. Abnormal vasculature was defined as that in which appears in the tumour area but not the control hemisphere. The full extent of RN1-luc spread and infiltration was most visible in T2 weighted fast spin echo sequences rather than T1 or T2* scans post-contrast. Across multiple timepoints, in U87MG, the T2* signal was converted to bright contrast surrounding the tumour area, while for RN1-luc vascular signal was converted to positive contrast, though the full extent of RN1-luc was still not entirely visible using this method (Figure 4, Figure S2).

**Figure 4:**
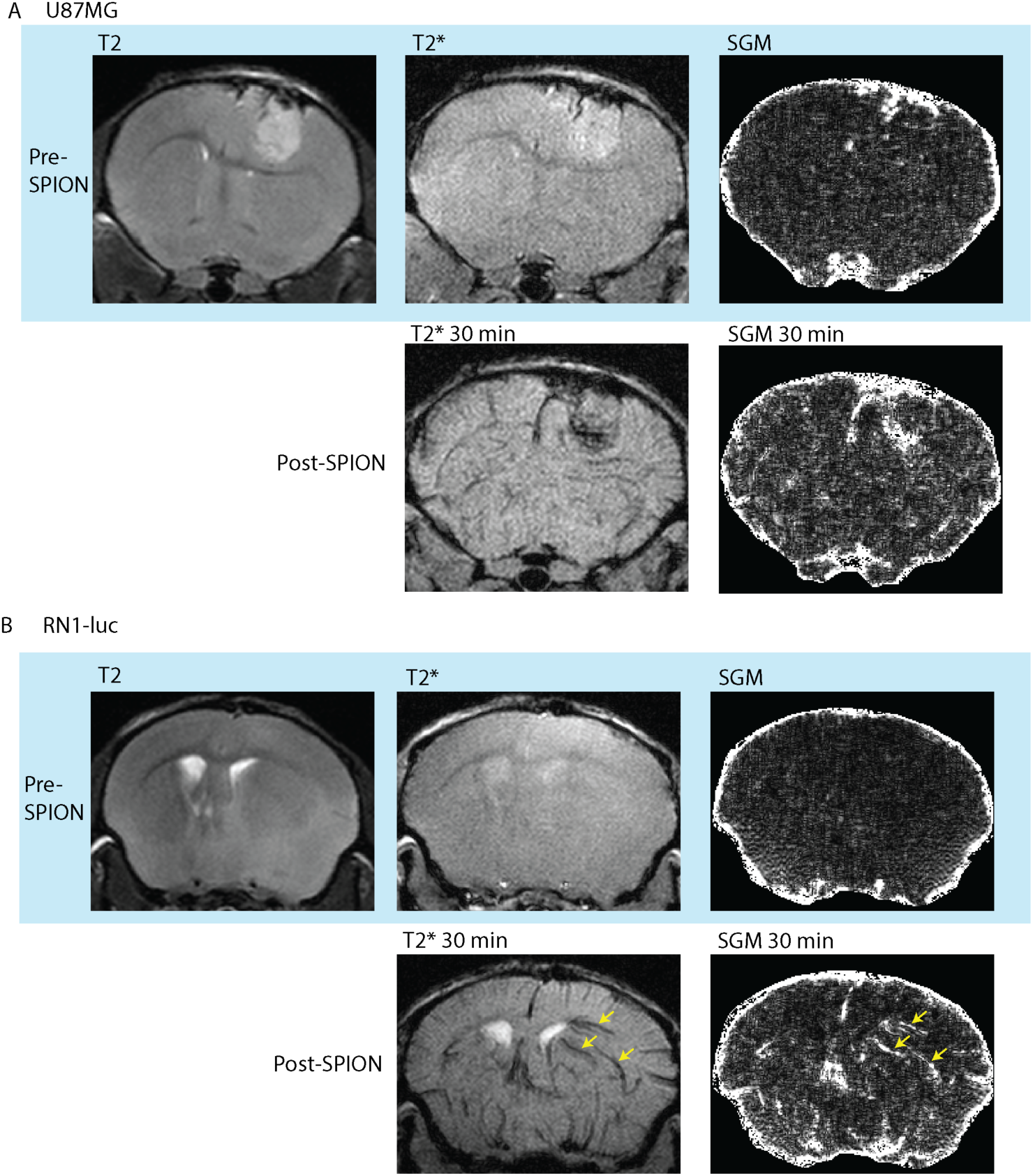
Bright contrast of PEG-SPIONs generated by susceptibility gradient mapping (SGM) post-processing of T2* weighted gradient recalled echo sequences calculated from equation 1, shown in **A)** U87MG and **B)** RN1-luc. In RN1-luc, abnormal blood vessels are visible in the right hemisphere post-SPION injection (yellow arrows). U87MG, which is a more permeable tumour, shows a high concentration of PEG-SPION in a highly vascularised region.

Figure S5 indicates a large susceptibility gradient at the kidneys, stomach, liver, and spleen 1 h post PEG-SPION injection, indicating interfaces between iron-rich areas and the surrounding iron-poor tissue. The insides of these organs remain dark, indicating a homogeneous distribution of nanoparticles within the organ, which in turn results in a lack of large susceptibility gradient vectors within the organ and thus little-to-no bright contrast is seen. In organs with a very high concentration of SPIONs, there is no measurable signal in these regions due to the short T2*. Hence, signal voids are produced in the image and as a result no SGM can be calculated for these regions. This serves in contrast to hypointensities in the lung, which are dark both pre- and post-injection and are not as significantly highlighted by this post-processing method – due to the presence of air and a low signal-to-noise ratio which prevents accurate calculation of the susceptibility gradient.

### Biodistribution results

A comparison of T2* weighted MR images pre-injection and 1 hour post PEG-SPION injection (Figure 5) revealed a decrease in the MGV post-injection in the liver and spleen. A similar biodistribution profile was observed for ferumoxytol (Figure S3). The strong inhomogeneity in magnetic susceptibility caused by the SPIONs in vasculature also appeared to enhance contrast within the kidneys, with the renal vasculature being sharply defined against the surrounding renal medulla, but only at early time points (generally <1 h) post injection (Figure 5, Figure S4).

**Figure 5:**
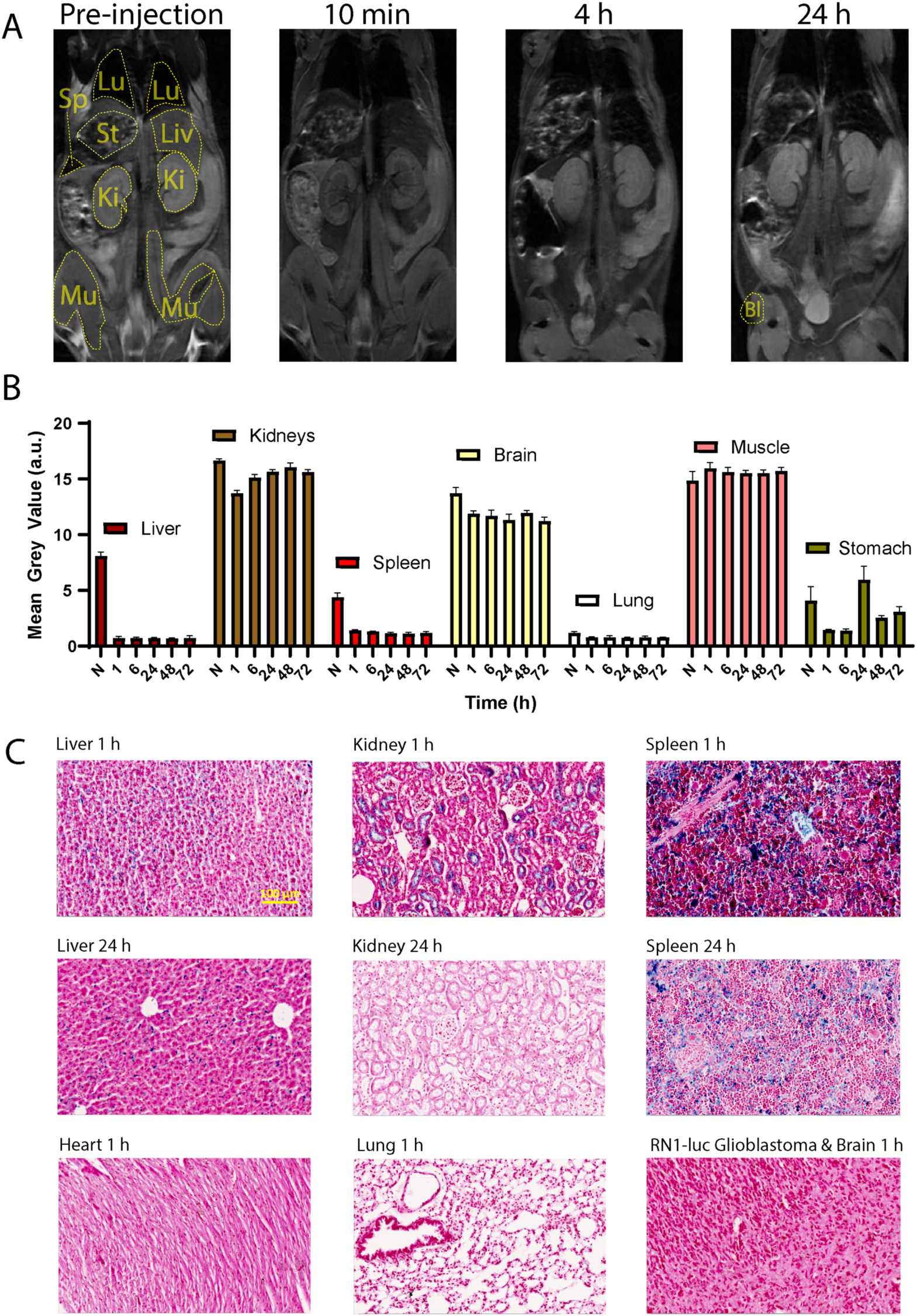
**A)** Biodistribution of the PEG-SPIONs (representative image, n=3), as shown by 7T MR T1-weighted spin echo images in NOD-SCID mice. From left to right, top to bottom, lung (Lu), stomach (St), liver (Li), spleen (Sp), kidney (Ki), muscles (Mu) and bladder (Bl) are encircled in yellow. Displayed window width and window level are equal for all images. **B)** Quantification of unscaled reconstructed k-space of 3 T T2*-weighted gradient recalled echo acquisitions in C57BL/6 mice over longer time points. Due to the proportions of the body coils, brains were not visible in 7 T body images. Graph represents mean + SEM grey values (arbitrary units) from oval-shaped regions of interest in T2* weighted FLASH 3 T images (n=2). **C)** Representative histology images of 10 micron sections stained with Perls’ Prussian blue and 0.1% safranin O. Iron oxide is shown by blue colour, notably in the liver and spleen, and to some extent in the kidney at 1 h. Scale bar = 100 microns.

There was little overall difference between quantification of images in terms of SPIONs T2 susceptibility effects in 3 T versus 7 T scanners, nor were there any strong differences in the peripheral organ changes between T1, T2 and T2* weighted images (data not shown). Uptake of the nanoparticles into the liver and spleen parenchyma was confirmed using Perls’ Prussian blue staining of 10 micron paraffin-embedded slices post transcardial saline and formalin perfusion (Figure 5).

## Discussion

Choice of orthotopic glioblastoma model is vital for evaluating nanoparticle, nanomedicine or other systemic therapeutic uptake and biodistribution^25^. The tumour border of U87MG is extremely clear in MRI scans as shown by both SPIONs and Gadovist contrast enhancement (Figure 3). RN1-luc tumours on the other hand, more faithfully recapitulate intratumoural BBB functionality, with the stem-like properties of the cells allowing growth as a diffuse dense mass within healthy tissue that spreads throughout the brain into healthy circuitry tissue^32^ over a longer time period. This infiltrative growth is difficult to clearly distinguish using MRI, though asymmetries and invasive changes in tissue density become evident in T2-weighted scans (Figure 2 and Figure 4). However, the true extent of the RN1-luc glioblastoma infiltration can only be visualised postmortem through histology and microscopy, which is also largely the case in human patients. Interestingly, Gadovist did not provide clear information regarding the infiltration extent of RN1-luc tumours, even in highly progressed cases (Figure 2), whereas the PEG-SPIONs provided information on abnormal vasculature and some angiogenic areas of the tumour (Figure 3, Figure 4, Figure S2).

Other radiological imaging modalities including positron emission tomography, magnetic resonance spectroscopy and diffusion tensor imaging may provide further qualitative information of glioblastoma location to that reported here. However, while the sensitivities of these modalities are favourable, the resolutions of these technologies are generally more limited than standard MRI sequences, and they have not yet been widely incorporated into clinical workflows. Two limitations for using diffuse glioblastoma patient-derived xenografts in our hands, are the long *in vivo* growth period of over 90 or 155 days, and the compromised immune system required. This long growth period constrains the time efficiency of clinically relevant preclinical studies. Furthermore, any passage of patient derived xenografts as allografts to maintain tumourigenicity inevitably changes the phenotype of the cells compared to the original host tumour^24,35^. Use of a genetic glioblastoma models such as triple PTEN, Trp53, and Rb1 mutation induction via a Cre-loxP system also has a long tumour growth period of approximately 100-200 days, and though these mice have a functional immune system, the growth location of each individual tumour is unpredictable^36^.

The PEGylated nanoparticles used in this study exhibit strong MRI vascular contrast at doses compatible with clinical use, and appear to be phagocytosed by hepatic and splenic reticuloendothelial macrophages of the mononuclear phagocytic system (Figure 5), following a several-hour blood circulation and blood-pool T2* contrast effects. Similar to clinically used carbohydrate-coated SPIONs such as ferumoxytol (Feraheme®), PEGylated SPIONs may have applications in not only glioblastoma angiogenesis MRI, but also possibly as a general alternative to gadolinium-based contrast. Careful modification of the surface coating of SPION contrast agents with high affinity targeting ligands may offer further highlighting of aggressive tumour infiltration. Future studies may explore how antibody- or ligand-targeted and passive PEGylated stealth nanoparticles are clinically applicable to glioblastoma prognosis and treatment response monitoring and evaluation. Feasibility of large-scale Good Manufacturing Practice grade nanoparticle production will also be an important consideration for future clinical implementation.

SPIONs such as ferumoxytol are prescribed off-label for clinical neuroimaging purposes at concentrations of ~1-11 mg Fe/kg (up to a 510 mg bolus)^37^, corresponding well in the range of that used in this study. Considering differences in allometry between mice and humans, the human equivalent dose for 10 mg/kg in mice is approximately 0.8 mg/kg^38^. The optimal concentration for imaging is governed by the iron-oxide contrast agent pharmacological, metabolic, and physical properties. A key physical property is the relaxivity of the contrast agent at the field strength chosen for MR imaging. The nanoparticles used here were initially developed for use as a contrast agent in superparamagnetic relaxometry (magnetic particle imaging, MPI) and were thus tuned for very high saturation magnetization that gives high transverse relaxivity (r2 = 177 ± 9 mM^−1^ s^−1^ at 3 T), while the r1 of this nanoparticle on the other hand is close to 0 at clinical field strengths ^30^. Therefore, this nanoparticle is a strong candidate for use as a MRI contrast agent, as its saturation r2 relaxivity is approximately 2.8 times greater than that of clinically used ferumoxytol^35^ (r2 = 62.3 ± 3.7 s^−1^ mM^−1^) and approximately 3.33 that of ferumoxtran-10 (r2 = 53.1 ± 3.3 s^−1^ mM^−1^)^36^. In principle this allows for the administration of a smaller quantity of PEG-SPION while maintaining still-substantive T2 dark contrast in regions of low SPION uptake, where contrast enhancement with existing clinical methods would not be possible. This passive PEG-SPION may therefore have also strong applications in functional liver MRI (Figure 5), as the liver rapidly uptakes SPIONs with little evidence of clearance across a 72-hour period, while non-functional liver would be expected to show a lack of uptake^37^. Hepatic Kupffer-Browicz cells are known to be highly endocytic and readily phagocytose most nanoparticles, particularly iron oxide nanoparticles^39,40^. Furthermore, with the advent of affordable, portable MRIs reaching the market, such as the Hyperfine Swoop® system, future low-cost screening of early-stage cancers such as in breast, prostate, liver and brain using affordable MR is becoming increasingly viable, and in such cases low-toxicity, high-sensitivity contrast agents are paramount.

There are well over 100 preclinical nanoparticle studies published in recent years for the magnetic resonance imaging of glioblastoma^41^. However, a large proportion of these rely solely on U87MG tumours^25,41^ or similar differentiated C6 or 9L tumours, and in some cases flank subcutaneous xenografts which all have poorly relevant blood brain barrier properties. Ferumoxtran-10 and ferumoxytol have been imaged in U87MG and human glioblastoma cases for over 20 years, with strong uptake appearing at approximately 24 hours post-injection^42–44^. However, the engulfment of these iron oxide nanoparticles appears to be largely by reactive macrophages rather than the tumour cells themselves^44^, which is different to what we observed regarding the wash in and wash out signal of PEG-SPION in U87MG and RN1-luc. Recent clinical studies comparing ferumoxytol to GBCAs have found differences in glioblastoma borders and dynamic susceptibility contrast, further indicating that these dextran-coated SPIONs are predominantly engulfed by tumour associated macrophages^43,45^, though by the lack of iron staining in brain tumour tissue post-perfusion in our study, it is doubtful that this is the case for PEG-coated SPIONs.

Our biodistribution results are consistent with those of Gobbo *et al.* ^46^, who administered mice with 12-15 nm SPIONs coated with dimercaptosuccinic acid at an intravenous concentration of 10 mg Fe/kg. T2-weighted MR imaging was performed at 7 T using turbo spin echo sequences post intravenous injection. They observed a maximum change in MRI signal intensity at the 3-hour mark for both the liver and spleen, from which the signal remained relatively constant up-unto the 96-hour mark, and a blood half-life of 32 min. In our study, we found rapid uptake of the PEG-SPION by the liver, spleen and less so in the kidneys within 10 min to 1 hr. For the kidneys in Gobbo *et al.*^46^, a signal peak was reached 48 hours post-injection, with a gradual decline observed out until the 96-hour mark. This is indicative of a slower renal uptake and clearance of the dimercaptosuccinic acid coated 15 nm SPIONs when compared to PEG-SPION.

In another study, Pham *et al*.^47^ obtained *ex vivo* biodistribution and clearance profiles in mice intraperitoneally-administered novel 25 nm core, methoxypolyethylene glycol coated SPIONs. This analysis quantified the iron concentrations in various peripheral organs via atomic absorption spectroscopy measurements. The mice were administered with a much larger dose of SPIONs – 90 mg Fe/kg – and were examined up to 1-week post-injection. Iron concentrations in the liver, spleen, kidney, and bladder all peaked at 1 hour post injection, with observed iron concentrations being relatively stable up to 1-week post injection, aside from a gradual reduction in iron content of the kidney by the 48-hour mark. In our *in vivo* imaging study, we observed a more rapid clearance of nanoparticles from both the kidney (Figures 5) and bladder (data not shown) by the 24-hour mark, which may also be related to differences between intravenous and intraperitoneal administration.

The SGM postprocessing step was performed to create image hypointensities as a result of SPION presence in the body, as regions of high SPION uptake will produce a strong susceptibility gradient at interfaces with regions of low SPION uptake. Positive contrast from iron-oxide nanoparticles can also be achieved via sequences such as balanced steady-state free precession, though this would be challenging at the field strengths used here due to banding artefacts^30,31^. The benefits of this SGM post-processing approach when compared to other approaches for generating bright contrast from dark T2 SPION MR contrast include a higher sensitivity to field inhomogeneity and susceptibility gradients combined with a reasonable suppression of background signal and water signals. Additionally, the signal-to-noise ratio is high when compared to other methods^26,48,49^.

The sensitivity to local susceptibility that the SGM approach provides in tandem with the high magnetization of the nanoparticle may have applications in the detection of microhemorrhages or calcification within tissue, as there will be a high susceptibility gradient between SPION-rich blood and these regions of interest^26,50^. As shown in Figure 4, this approach could have application in the detection of glioblastoma, where SPIONs continue to be investigated for their applicability as an alternative to gadolinium-based MR contrast agent due to possible toxicity from gadolinium deposition, especially in patients with multiple administrations^51^.

In conclusion, PEGylated SPIONs provide additional vasculature information to that obtained with conventional glioblastoma MRI. While small gadolinium-based contrast agents and ultrasmall SPIONs have shown rapid renal uptake^52^ and deposition throughout the body^53^, medium-sized SPIONs such as those used in this study appear to show a more preferential hepatobiliary macrophagic (Kupffer-Browicz cell) and splenic uptake, where it predominantly remains and is slowly metabolised. Further investigations in humans may be made using SGM approaches to test the clinical utility of positive contrast imaging of SPIONs.

## Acknowledgments

We acknowledge the support of the Sydney Imaging and National Imaging Facility, a National Collaborative Research Infrastructure Strategy (NCRIS) capability at Charles Perkins Centre, including Dr Nguyen Pham, Dr Sofie Trajanovska and Dr Raj Parajuli, for assistance with the MR imaging. We also thank Professor Bryan Day and QCell for providing the RN1-luc patient-derived xenograft cells, Dr Kelly McKelvey for providing advice on animal surgeries, and the University of Sydney Laboratory Animal Services including the Kearns Facility for housing of the mice. We acknowledge the use of services provided by the Preclinical pipeline for Advanced Cancer Therapeutics (PACT) at the Kolling Institute. We thank Imagion Biosciences for providing the nanoparticles used in this study, Dr Emily Hewson and Jeremy Lim for providing critical reading of the manuscript, Dr Samson Dowland for histological advice and Dr Yingying Su for assistance with microscopy. PWJ was supported by a Cancer Institute New South Wales Translational Program Grant (TPG2165). DEJW was supported by Australian National Health and Medical Research Council (NHMRC) Investigator Grant 2017140. CB and TB were supported by NHMRC Investigator Grant 1194004.

## Author Contributions

The study was conceptualized by D.E.J.W., C.B., Z.K., and P.W.J. Methodology was developed by P.W.J., D.E.J.W., R.M., Y.G., C.B., E.G.K., B.S., M.Z., S.S., T.D.T.T., F.W., and A.T. MRI investigations were conducted by P.W.J., R.M., and Y.G. Data visualization and analysis were handled by P.W.J., R.M., T.B., and D.E.J.W. Funding was secured by Z.K. and D.E.J.W. Project administration was managed by D.E.J.W. and Z.K. Supervision was provided by D.E.J.W., Z.K., and L.M. The original draft was written by P.W.J. and D.E.J.W. All authors contributed to review and editing of the final manuscript.

## Data Availability

The authors declare that original data supporting the findings of this research are available within the article. Raw data is available from the corresponding author upon reasonable request.

## Abbreviations

BBB: blood-brain barrier
FA: flip angle
FOV: field of view
GRE: gradient recalled echo
MGV: mean grey value
MRI: magnetic resonance imaging
NA: number of averages
PDX: patient-derived xenograft
PEG: polyethylene glycol
T1w: T1-weighted
T2w: T2-weighted
T2*w: T2*-weighted
TE: echo time
TR: repetition time
SGM: susceptibility gradient mapping
ST: slice thickness
SPION: superparamagnetic iron oxide nanoparticle

## Supplementary Figures

**Figure S1 –.**
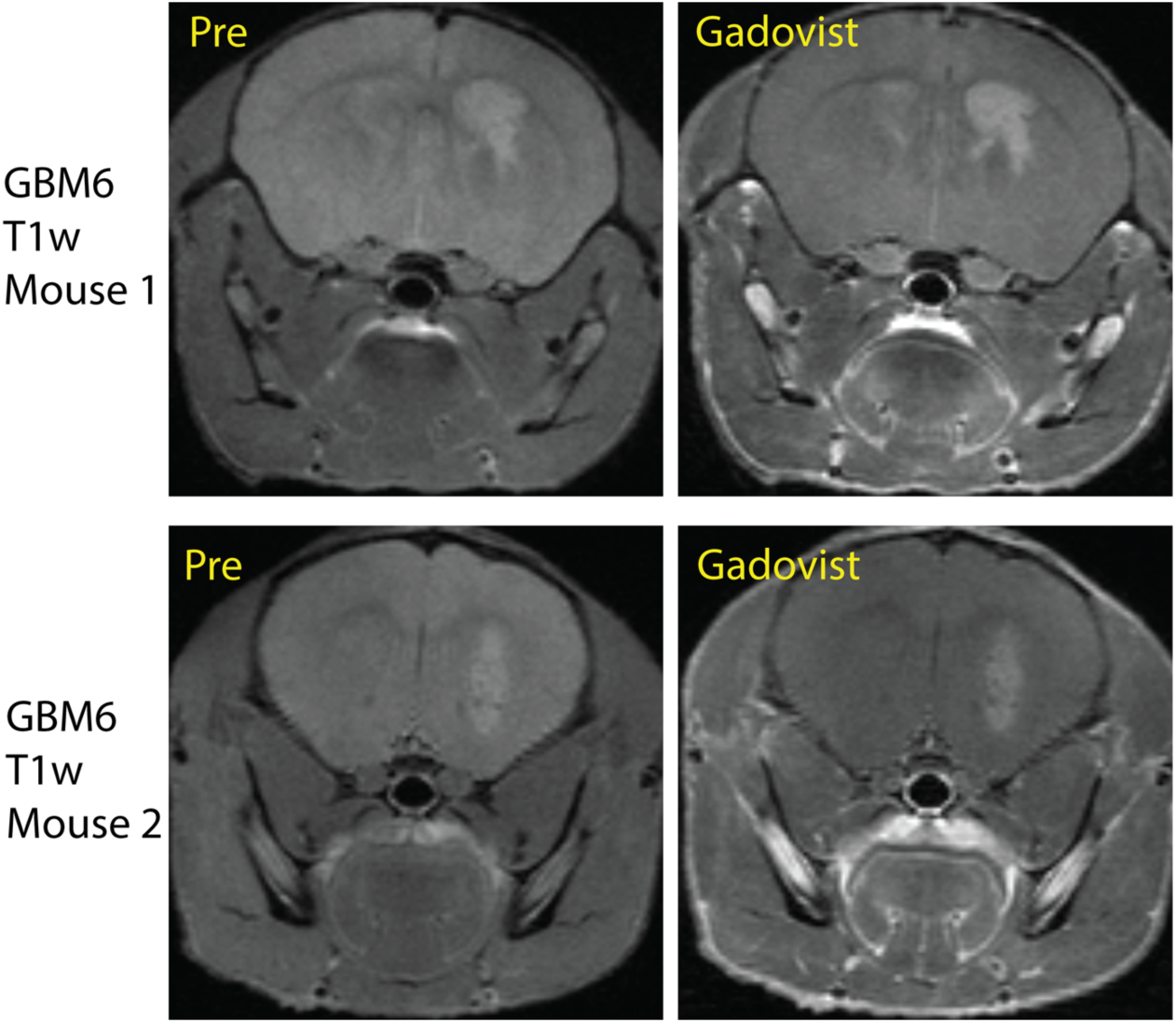
Two examples of patient derived xenograft GBM6 pre- and post-Gadovist (0.1 mmol/kg), as shown in T1-weighted turbo spin echo scans. Permeability of the tumour BBB appears limited at this stage of growth due to modest Gadovist (Gadolinium) enhancement.

**Figure S2:**
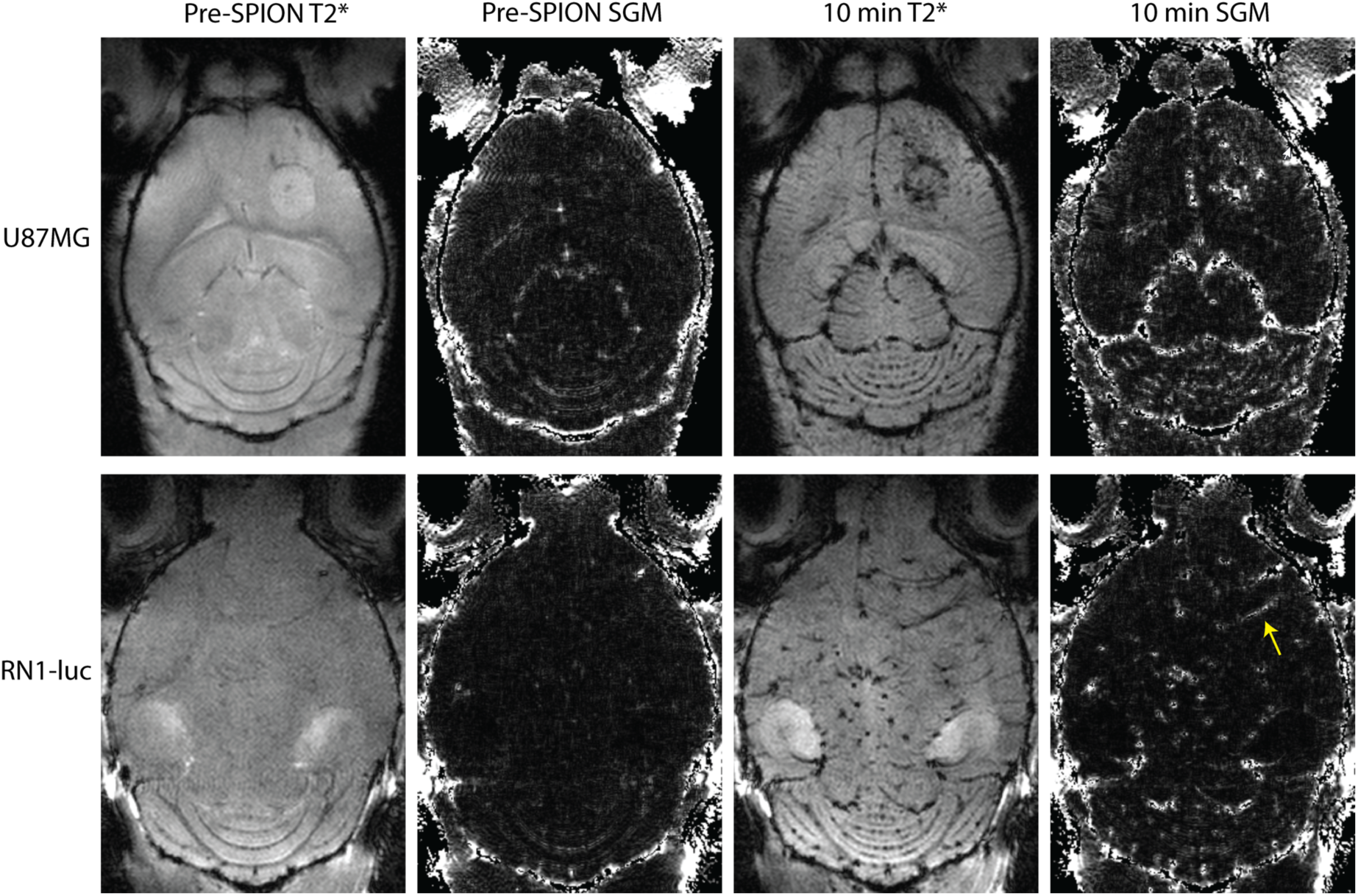
Additional susceptibility gradient mapping (SGM) images adjacent to source T2* gradient recalled echo sequences in U87MG and RN1-luc glioblastoma xenografts pre and post-10 mg/kg PEG-SPION from a horizontal field of view. Yellow arrow points to an example of diffuse glioblastoma angiogenesis as shown by PEG-SPION positive contrast in SGM.

**Figure S3:**
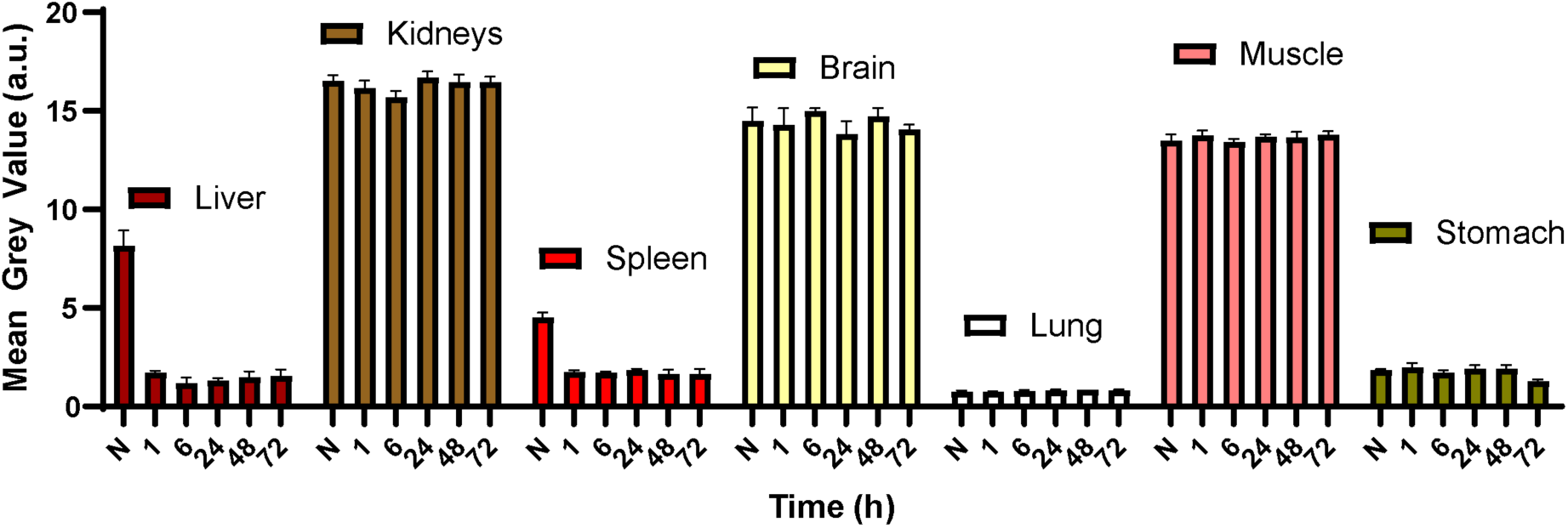
Mean + SEM pharmacokinetic biodistribution of 2 mg/kg ferumoxytol from T2*-weighted gradient recalled echo sequences (biological n=1, number of ROIs across slices per timepoint = 2 to 8).

**Figure S4:**
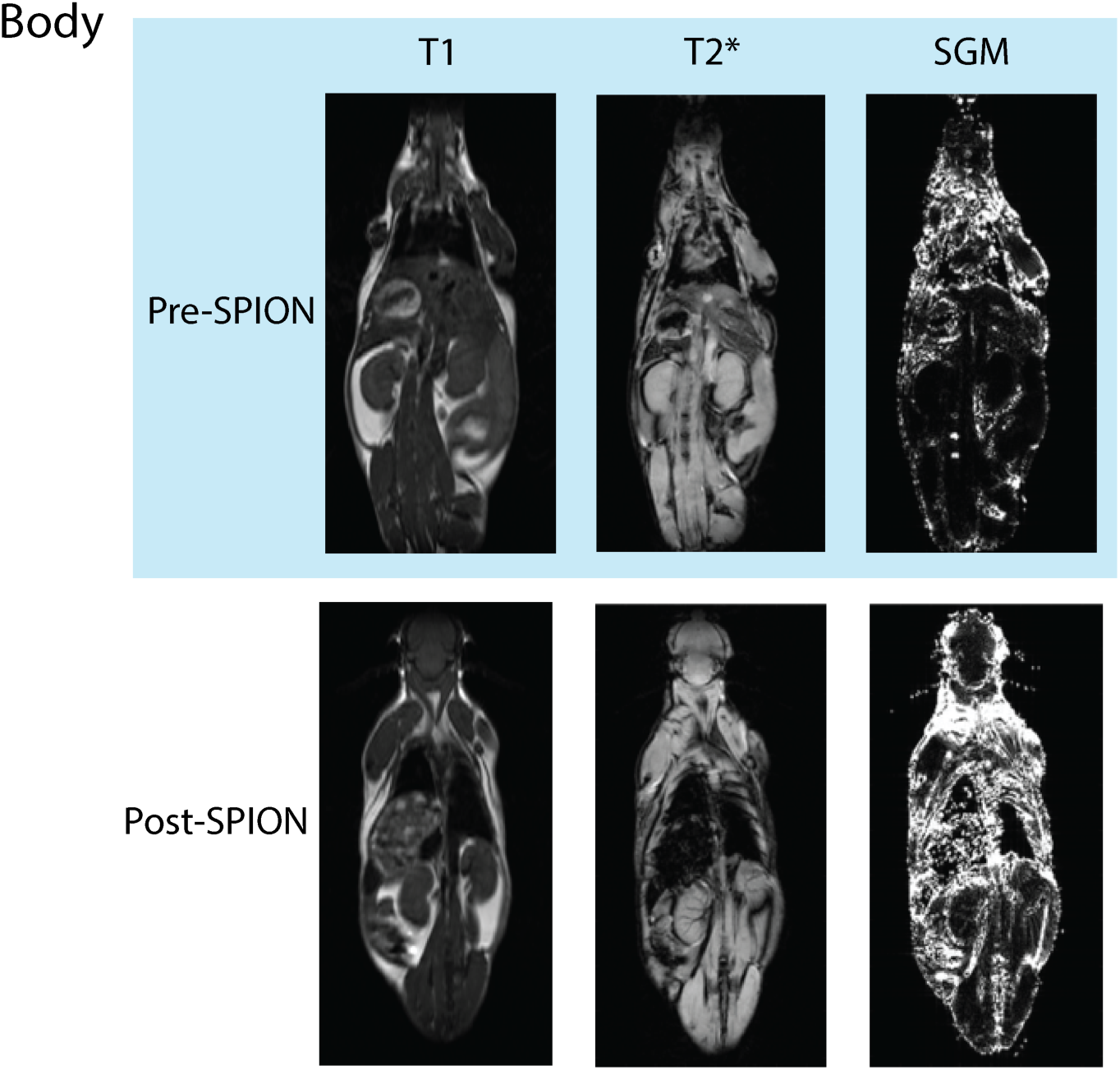
Body 3 T T1 weighted and T2* weighted scans, plus susceptibility gradient maps pre and 60 minutes post 10 mg/kg PEG-SPION injection.

